# Differential effects of voclosporin and tacrolimus on insulin secretion from human islets

**DOI:** 10.1101/2020.04.30.071324

**Authors:** Jelena Kolic, Leanne Beet, Peter Overby, Haoning Howard Cen, Evgeniy Panzhinskiy, Daren R. Ure, Jennifer L. Cross, Robert B. Huizinga, James D. Johnson

## Abstract

**Context:** The incidence of new onset diabetes after transplant (NODAT) has increased over the past decade, likely due to calcineurin inhibitor-based immunosuppressants, including tacrolimus (TAC) and cyclosporin (CsA). Voclosporin (VCS), a next generation calcineurin inhibitor is reported to cause fewer incidences of NODAT but the reason is unclear.

**Objective:** Whilst calcineurin signaling plays important roles in pancreatic β-cell survival, proliferation, and function, its effects on human β-cells remain understudied. In particular, we do not understand why some calcineurin inhibitors have more profound effects on the incidence of NODAT.

**Methods:** We compared the effects of TAC and VCS on the dynamics of insulin secretory function, programmed cell death rate, and the transcriptomic profile of human islets. We studied two clinically relevant doses of TAC (10 ng/ml, 30 ng/ml) and VCS (20 ng/ml, 60 ng/ml), meant to approximate the clinical trough and peak concentrations.

**Results:** TAC, but not VCS, caused a significant impairment of 15 mM glucose-stimulated and 30 mM KCl-stimulated insulin secretion. This points to molecular defects in the distal stages of exocytosis after voltage-gated Ca^2+^ entry. No significant effects on islet cell survival or total insulin content were identified. RNA sequencing showed that TAC significantly decreased the expression of 17 genes, including direct and indirect regulators of exocytosis (*SYT16*, *TBC1D30*, *PCK1*, *SMOC1*, *SYT5, PDK4*, and *CREM*), whereas VCS has less broad and milder effects on gene expression.

**Conclusions:** Clinically relevant doses of TAC, but not VCS, directly inhibit insulin secretion from human islets, likely via transcriptional control of exocytosis machinery.

## Introduction

New onset diabetes after transplant (NODAT) is a clinical problem that has increased in incidence over the past decade. It is widely believed that NODAT is caused by exposure to steroids or high dose calcineurin inhibitors, including tacrolimus (TAC; also known as FK506) and cyclosporin (CsA) (1,2). These drugs are shown to damage cell types critical for the maintenance of glucose homeostasis, particularly pancreatic β-cells (3). TAC is a mainstay for prevention of transplant rejection, but effective immunosuppression targeting a concentration range of 4-11 ng/mL still results in NODAT in > 20% of patients (1). There is an unmet clinical need for calcineurin inhibitors that do not cause diabetes in this already vulnerable patient population.

Voclosporin (VCS) is a next-generation calcineurin inhibitor that is structurally related to cyclosporine A (CsA) with the addition of a carbon molecule at amino acid-1 of CsA. This modification results in enhanced binding of the VCS-cyclophilin complex to calcineurin and shifts metabolism, resulting in increased potency and a consistently better pharmacokinetic-pharmacodynamic profile as compared to CsA. VCS was previously evaluated in a Phase IIb, multi-center, open-label, concentration-controlled study in patients undergoing *de novo* renal transplantation, evaluating the efficacy and safety of three doses of VCS (0.4, 0.6, and 0.8 mg/kg BID) as compared to TAC (4). In that study, the low-dose VCS group had similar efficacy to tacrolimus in controlling acute rejection with a significantly decreased incidence of NODAT (1.6% vs. 16.4%, respectively, p=0.031) (4). In the AURA study for lupus nephritis, 1.1% of patients receiving VCS (23.7 mg) and 1.1% of patients in the placebo group reported diabetes (5), contrasting with NODAT in 25.7% of patients while taking a standard dose of TAC for 1 year (6). Thus, there is an unexplained difference between the clinical diabetes seen after treatment with VCS, and currently prescribed immunosuppressants, TAC and CsA.

While the etiology of NODAT is not well understood, calcineurin and nuclear factor of activated T-cells (NFAT) are involved in a number of cellular processes outside of immunosuppression, including direct roles in normal pancreatic β-cell development and physiology (7–9). For example, Heit et al. reported that β-cell selective knockout of calcineurin in mice reduced functional β-cell mass and caused diabetes that could be rescued by over-expressing active NFATc1 (8). The effects of acute calcineurin inhibition on human and mouse islet function was previously investigated in a study by our group comparing clinically relevant concentrations of TAC, CsA, and rapamycin (7). *In vitro* experiments showed a direct decrease in β-cell function after only 24 hours of exposure to TAC (7). *In vivo* studies in mice from our group and others have shown that inhibition of the calcineurin pathway by TAC impairs the function of transplanted human islet grafts (7,10). Thus, while previous data from animal models and human islets studies clearly demonstrates that TAC has multiple pathological effects on pancreatic islets, the mechanisms by which TAC impairs insulin secretion from human islets remained unresolved. Most importantly, there are no published studies directly comparing the direct effects of TAC versus VCS on human islets.

Here, we directly tested the hypothesis that VCS has lower human β-cell toxicity compared with TAC, and we conducted studies designed to investigate the molecular mechanisms involved (Fig 1A). We found that human islets exposed to TAC exhibited significantly reduced distal steps of glucose-stimulated insulin secretion, while VCS showed no statistically significant inhibition at a dose that elicits sufficient immunosuppression. RNA sequencing showed that TAC, and to a lesser extent VCS, decreased the expression of genes that specifically regulate the distal steps of insulin secretion. In support of the clinical observations, our data suggest that VCS is less toxic to human pancreatic β-cells at clinically relevant doses.

**Figure 1.**
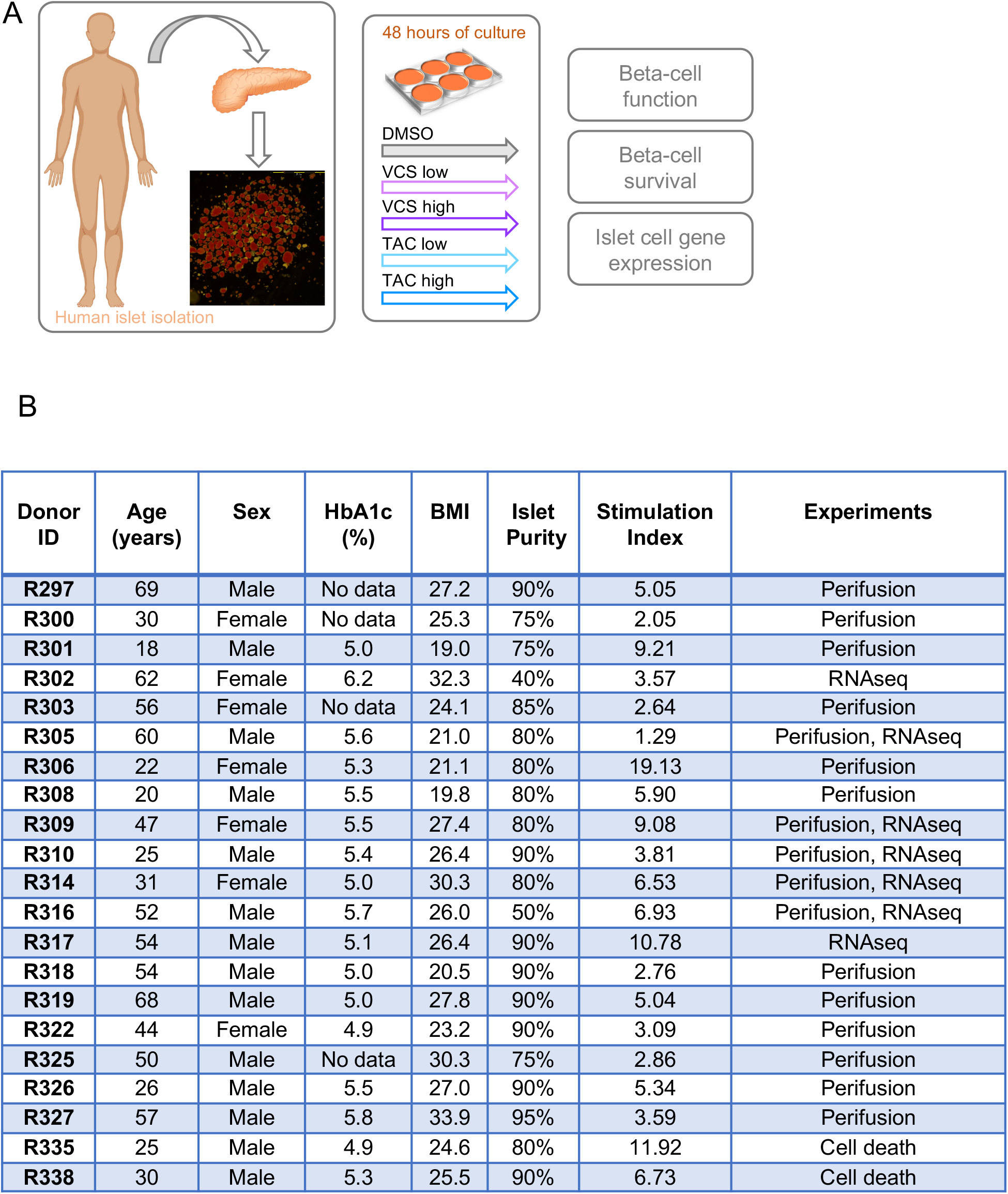
Experimental design, and human islet donor and isolation characteristics. **(A)** Experimental design for study of VCS and TAC effects on high quality primary human islets isolated from cadaveric donors. Details of each isolation are available from IsletCore.ca (the example image shown from the R297 isolation). **(B)** High-quality, research-specific human islets from consenting donors were obtained from IsletCore Laboratory in Edmonton, Alberta. Islet donors and isolation characteristics are listed here. Stimulation index is the average of 3 technical replicates of the 16.7 mM to 2.8 mM glucose response as provided by the IsletCore Laboratory quality control database.

## Methods

### Human islet culture

High-quality research specific human islets from cadaveric donors were obtained from IsletCore Laboratory in Edmonton Alberta (11), and used under ethical approval from the UBC Clinical Research Ethics Board (H13-01865). Islets were shipped in CMRL media (Thermo Fisher Scientific) overnight from Edmonton to the University of British Columbia. Upon arrival, islets were immediately purified by handpicking under a stereomicroscope and suspended in RPMI 1640 medium (Thermo Fisher Scientific) supplemented with 5.5 mmol glucose, 10% FBS, and 100 units/ml penicillin/streptomycin. Groups of 100 islets were placed in non-treated 100-mm polystyrene petri dishes (Fisher Scientific) followed by 48-hr incubation at 37°C and 5% CO_2_ (Fig. 1A). We chose 48-hr as an incubation time to balance our efforts to model long-term (years) exposure to these drugs in the clinical setting with our experience that human islet function degrades after 4-5 days in culture. Donor and isolation characteristics can be found in Figure 1B.

### Immunosuppression-related assays

The Calcineurin inhibitor assay and the T-cell proliferation assay are described in more detail elsewhere (12,13). Briefly, peripheral blood mononuclear cell (PBMC) lysates were used in the calcineurin assay after 25 min stimulation with PMA-A23187. After lysis with a calcineurin assay buffer (no detergents) and 3 freeze-thaw cycles, the soluble fraction was probed by Western blot with 1:200 anti-NFAT2 antibody (Santa Cruz Biotechnology Cat# sc-13033, RRID:AB_2152501). Western blot analysis of NFAT accumulation after treatment with indicated doses of VCS and TAC was then conducted to identify the relative concentrations that elicited a similar response.

### Dynamic analysis of insulin secretion from human islets

To define the dynamic effects of VCS and TAC on insulin secretion from human islets, we used human islet perifusion studies (7). We assessed the effects of TAC (10 ng/ml, 30 ng/ml) and VCS (20 ng/ml, 60 ng/ml) on the dynamics of insulin secretion in response to two stimuli that are diagnostic of changes in metabolism/signalling or exocytosis. Our standard approach (7,14–19) compared the response to 15 mM glucose stimulation and direct depolarization with 30 mM KCl. More specifically, 65 islets per column were perifused (0.4 ml/min) with 3 mM glucose KRB solution as described previously (7) for 60 min to equilibrate the islets to the KRB and flow rate, and then with the indicated condition (as described in corresponding figure legends). First phase insulin release was defined as the amount of insulin secreted during the first 20 min of 15 mM glucose stimulation, while the remaining 40 of stimulation were defined as second phase. DMSO, VCS and TAC were present at the indicated concentrations during the entirety of the perifusion experiment. We repeated this experiment on islets from 17 donors to take into account the significant variability between human islet preparations (11). Samples were stored at −20 °C and insulin secretion was quantified using human insulin radioimmunoassay kits (Cedarlane).

### Cell death assays

Pancreatic islets from human cadaveric donors were dispersed and seeded into 384-well plates and cultured in similar manner as previously described (20–23). Cells were stained with 50 ng/mL Hoechst 33342 (Invitrogen) and 0.5 μg/mL propidium iodide (PI) (Sigma-Aldrich) in RPMI 1640 medium (Invitrogen) with 5 mM glucose (Sigma-Aldrich), 100 U/mL penicillin, 100 μg/mL streptomycin (Invitrogen), 10% vol/vol fetal bovine serum (FBS) (Invitrogen) with or without a cytokine cocktail (25 ng/mL TNF-α, 10 ng/mL IL-1β, and 10 ng/mL IFN-γ; R&D Systems). We have previously shown that neither of these chemicals affect islet cell viability on their own at these concentrations (23). After 2 hours of staining, cells were incubated with different concentrations of VCS or TAC (1, 10, 20, 30, 60, 120 ng/ml) or vehicle (DMSO) and with a cytokine cocktail (25 ng/mL TNF-α, 10 ng/mL IL-1β, and 10 ng/mL IFN-γ; R&D Systems). Immediately after the addition of the test drugs, the cells were imaged at 37°C and 5% CO_2_. Images were captured every second hour for 48 hours using a robotic microscope (ImageXpress^MICRO^ XLS, Molecular Devices) and analyzed using MetaXpress multi-wave cell score Software (Molecular Devices). To adjust for the difference in basal cell death for each individual run, each timepoint was normalized to the first timepoint then further normalized to the cytokine control.

### RNA-sequencing and pathway analysis

Groups of 200 islets from 7 independent donors were treated for 48 hours with vehicle (DMSO), TAC (30 ng/ml) or VCS (60 ng/ml) then immediately flash-frozen in liquid nitrogen and stored at - 80°C. RNA isolation, library preparation, RNA sequencing and bioinformatics support were provided by the UBC Biomedical Research Centre Core Facility. Briefly, sample quality control was performed using the Agilent 2100 Bioanalyzer. Qualifying samples were then prepped following the standard protocol for the NEBnext Ultra ii Stranded mRNA (New England Biolabs). Sequencing was performed on the Illumina NextSeq 500 with Paired End 42bp × 42bp reads. De-multiplexed read sequences were then aligned to the reference sequence using STAR aligners (STAR, RRID:SCR_015899) Assembly were estimated using Cufflinks (Cufflinks, RRID:SCR_014597) through bioinformatics apps available on Illumina Sequence Hub. Raw counts were analyzed by NetworkAnalyst 3.0 (www.networkanalyst.ca) (24). DeSeq package was selected for differential expression of the genes. Differentially expressed genes were identified as those with an adjusted *p* value < 0.05 using the comparison of TAC vs DMSO or VCS vs DMSO. To determine the relationship between the differentially expressed genes and the calcineurin-NFAT pathway, we extracted all 49 genes in the Gene Ontology Term “calcineurin-NFAT pathway” (accessed July 30, 2020 from http://www.informatics.jax.org/vocab/gene_ontology/GO:0033173). We removed genes from the calcineurin-NFAT list without human homologues and that were not expressed in our human islet RNAseq. We then combined the expressed calcineurin-NFAT list with the genes that were differentially expressed between DMSO and TAC and used the protein-protein network interaction analysis tools at string-db.org with the following setting (active interaction sources – all; medium confidence 0.4; max interactors 0).

### Statistics and data analysis

Data were analyzed using a one-way multiple comparisons ANOVA followed by a *post hoc* t-test using the Tukey HSD method, unless otherwise indicated. Data are expressed as means ± SEM, and p < 0.05 was considered significant. n values represent the number of unique islet donors studied.

## Results

### Effects of TAC and VCS on NFAT activity

Both TAC and VCS exert their immunosuppressive effects through inhibition of calcineurin phosphatase activity, preventing the dephosphorylation and activation of NFATc (25,26). Each of these two drugs is distinct with regard to its molecular mechanism of action. TAC interacts with the immunophilin FKBP12 prior to binding calcineurin, whereas VCS interacts with cyclophilins. The *in vitro* activity of TAC, VCS (and the structurally related inhibitor CsA) were compared in a calcineurin inhibition assay and for their downstream ability to inhibit the division of activated T cells. The IC_50_ of TAC against calcineurin was found to be approximately 4-fold lower than VCS, in accordance with clinical dosing where the therapeutic range of TAC is 5-30 ng/mL and VCS target trough concentrations are around 20-60 ng/mL (Fig 2A). Surprisingly, the potency difference between TAC and VCS widened to 50-fold, in favor of TAC, when looking at inhibition of T-cell proliferation (Fig 2A), leading to additional experiments comparing the impact of each drug on their common pathway effector, NFAT. Maximal activation of NFAT was observed at 56 ng/mL of VCS and only 1.4 ng/mL of TAC, a potency difference of 40-fold (Fig. 2B,C,D). These data demonstrate that, at therapeutic concentrations, TAC has a far more profound impact on NFAT than VCS (Fig. 2E).

**Figure 2.**
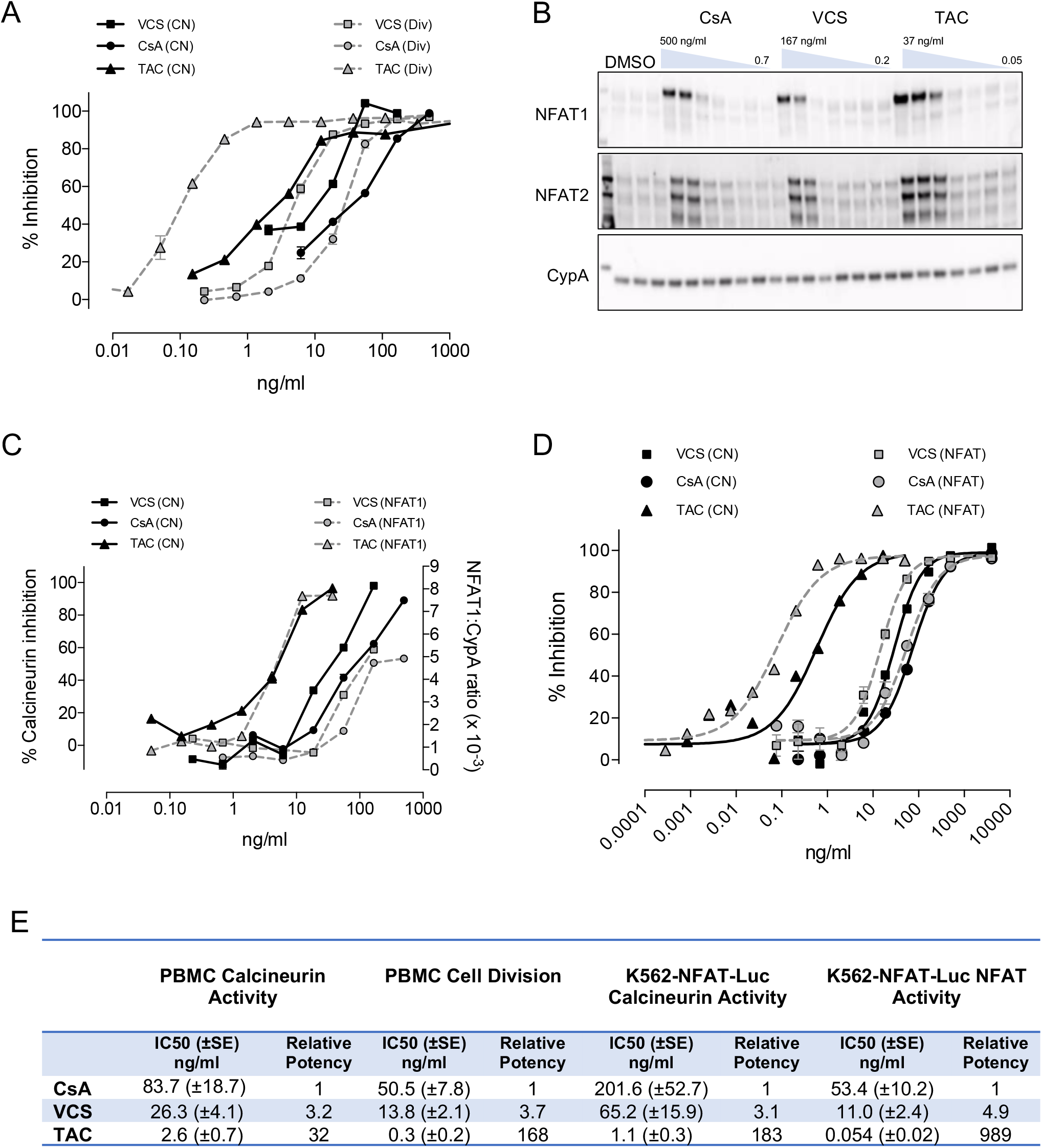
Differences between TAC, VCS and CsA on calcineurin activity, lymphocyte proliferation assay, and NFAT accumulation. **(A-D)** Sister aliquots of PBMC were treated in the presence of drug or vehicle control for analysis of lymphocyte cell division (Div), calcineurin activity (CN), and cytosolic NFAT. **(A)** Lymphocyte division was determined by CFSE analysis following 3-day stimulation with CD3 antibody. Calcineurin activity was determined by radioactive 32P-RII peptide assay following 30-min stimulation with PMA+A23187. **(B)** Cytosolic NFAT relative to CypA (loading control protein) was determined by Western blotting and densitometric quantitation following 30-min stimulation with PMA+A23187. **(C)** K562 cells stably transfected with an NFAT-luciferase reporter were stimulated for 22 hr with PMA + A23187 in the presence of drug or vehicle control. Cells were separated into separate aliquots for measurement of calcineurin activity and luciferase activity for NFAT promoter activation. **(D)** Percent inhibition was determined by normalizing values to vehicle control (100% activity) and to 2 μg/ml VCS (100% inhibition). These observations were replicated 5 times. **(E)** Calculated IC50 values in PBMC and K562-NFAT-Luc. n = 8-12 experiments for PBMC and 4-7 experiments for K562-NFAT-Luc.

### Effects of TAC or VCS on basal and glucose stimulated insulin secretion

Insulin is released from pancreatic islets, *in vivo* and *in vitro*, in a biphasic pattern that includes both a rapid 1^st^ phase of pre-docked insulin granules and a 2^nd^ sustained phase involving granules that are recruited to the plasma membrane (27). There is significant inter-individual variability in the rate of insulin secretion from human islets (11) and the effects of immunosuppressants (7), so we examined insulin secretion from high quality human islets isolated from 17 donors exclusively for research purposes. We employed the dynamic perifusion system with frequent sampling to evaluate basal insulin secretion at 3 mM glucose, both phases of insulin secretion during 15 mM glucose stimulation, and insulin secretion in response to direct depolarization with 30 mM KCl. TAC at 10 ng/ml (clinical trough) and 20 ng/ml (clinical peak) significantly inhibited total insulin secretion at 15 mM glucose, whereas VCS did not (Fig. 3A,B). TAC exhibited a specific effect on the 2^nd^ phase but not the 1^st^ phase of glucose-stimulated insulin secretion (Fig. 3C,D). The insulin secretory response to 30 mM KCl was also significantly inhibited by both concentrations of TAC (Fig. 3E). In contrast, VCS concentrations approximately 2 times higher than TAC had milder effects on glucose-stimulated insulin secretion. Specifically, the clinical trough concentration of 20 ng/ml VCS was ~50% less inhibitory to the total and 2^nd^ phase of insulin secretion and did not result in statistically significant inhibition (Fig. 3A,B,D). No significant differences in basal insulin secretion were observed at 3 mM glucose for either drug at either of the concentrations tested (Fig. 3F). Similar results were observed when the perifusion data were normalized to basal insulin secretion (Fig. 3G,H). We were unable to detect significant differences in acid-ethanol extractable insulin protein content in human islets treated with either drug, at either concentration, for 48 hr (Fig. 4). These results indicate that the specific significant inhibitory effects of TAC on total and 2^nd^ phase glucose-stimulated insulin secretion, as well as insulin secretion stimulated by direct depolarization, are likely to be direct effects on the exocytosis machinery rather than a result of reduced insulin production. Collectively, these data also demonstrate that VCS has milder effects on human islet secretory function when compared with TAC in this sensitive *in vitro* assay system.

**Figure 3.**
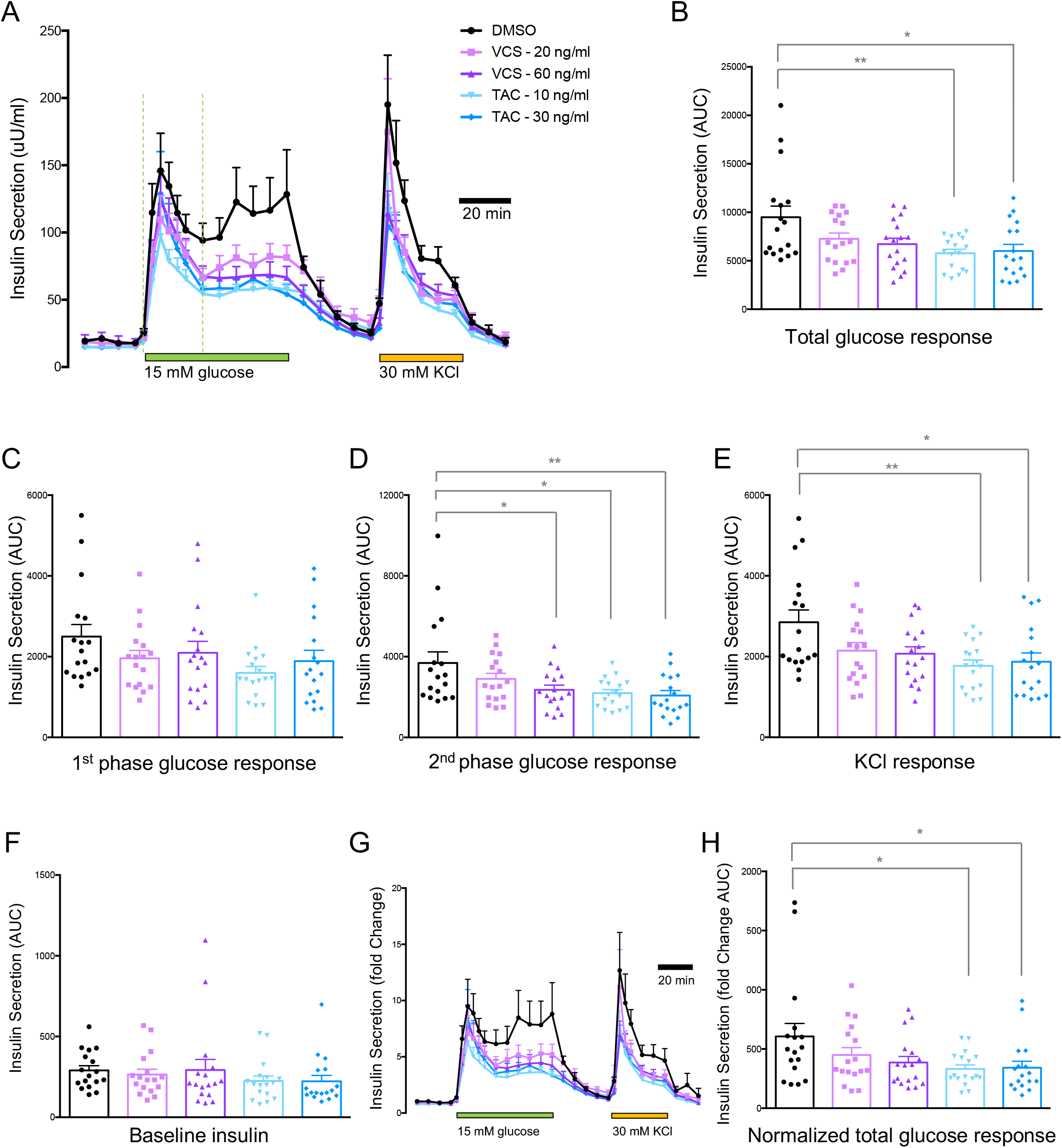
Basal, glucose-stimulated and KCl-stimulated insulin secretion from human islets treated with peak and trough concentrations of VCS and TAC. **(A)** Averaged traces of dynamic insulin secretion measurements in the context of 3 mM glucose, 15 mM glucose, or 30 mM KCl (as indicated). **(B)** Total area under the curve (AUC) of the 15 mM glucose response. (**C, D, E)** AUCs of 1^st^ phase and 2^nd^ phase 15 mM glucose responses, as well as the KCl response. **(F)** Baseline insulin secretion. **(G, H)** Insulin secretion normalized to baseline, including AUC. n=17

**Figure 4.**
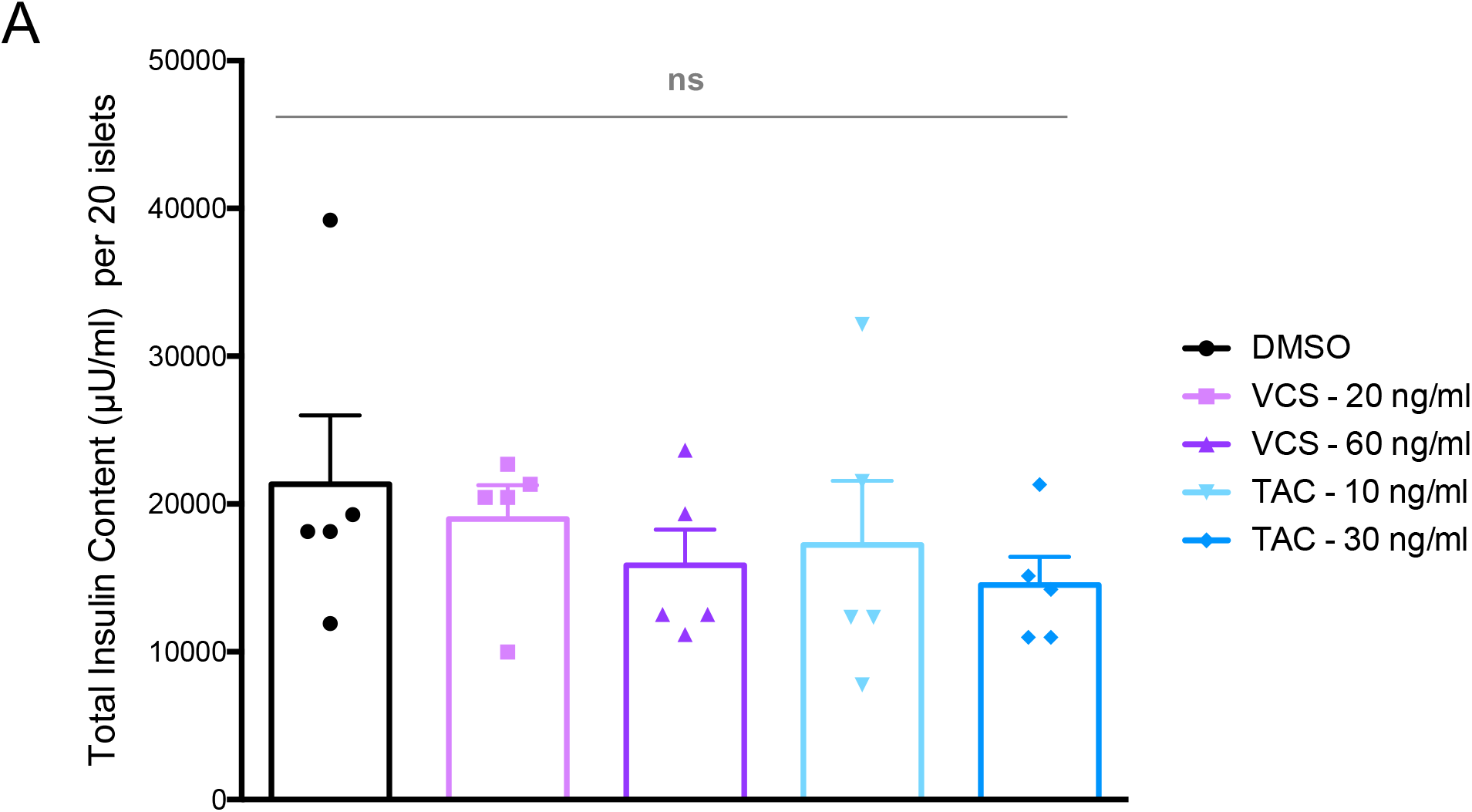
Insulin content from isolated human islets treated with VCS and TAC. Insulin protein measured by radioimmunoassay from 20 islets treated for 48 hours with VCS or TAC (concentrations as indicate) after acid-ethanol extraction of insulin.

### Effects of TAC or VCS on human islet cell survival

We next assessed the effects of TAC and VCS on the survival of dispersed human islet cells using our long-term high-content imaging protocols that measure both apoptotic and non-apoptotic cell death over multiple days of culture in the presence of proinflammatory cytokines (20–23) (Fig. 5A,B). We studied both TAC and VCS over a 5 point-dose response curve. We observed no significant differences in human islet cell survival following either drug treatment (Fig. 5C). This is consistent with our data showing no changes in insulin content following 48 hrs of drug treatment, further pointing to the primacy of the direct effect of TAC on insulin secretion.

**Figure 5.**
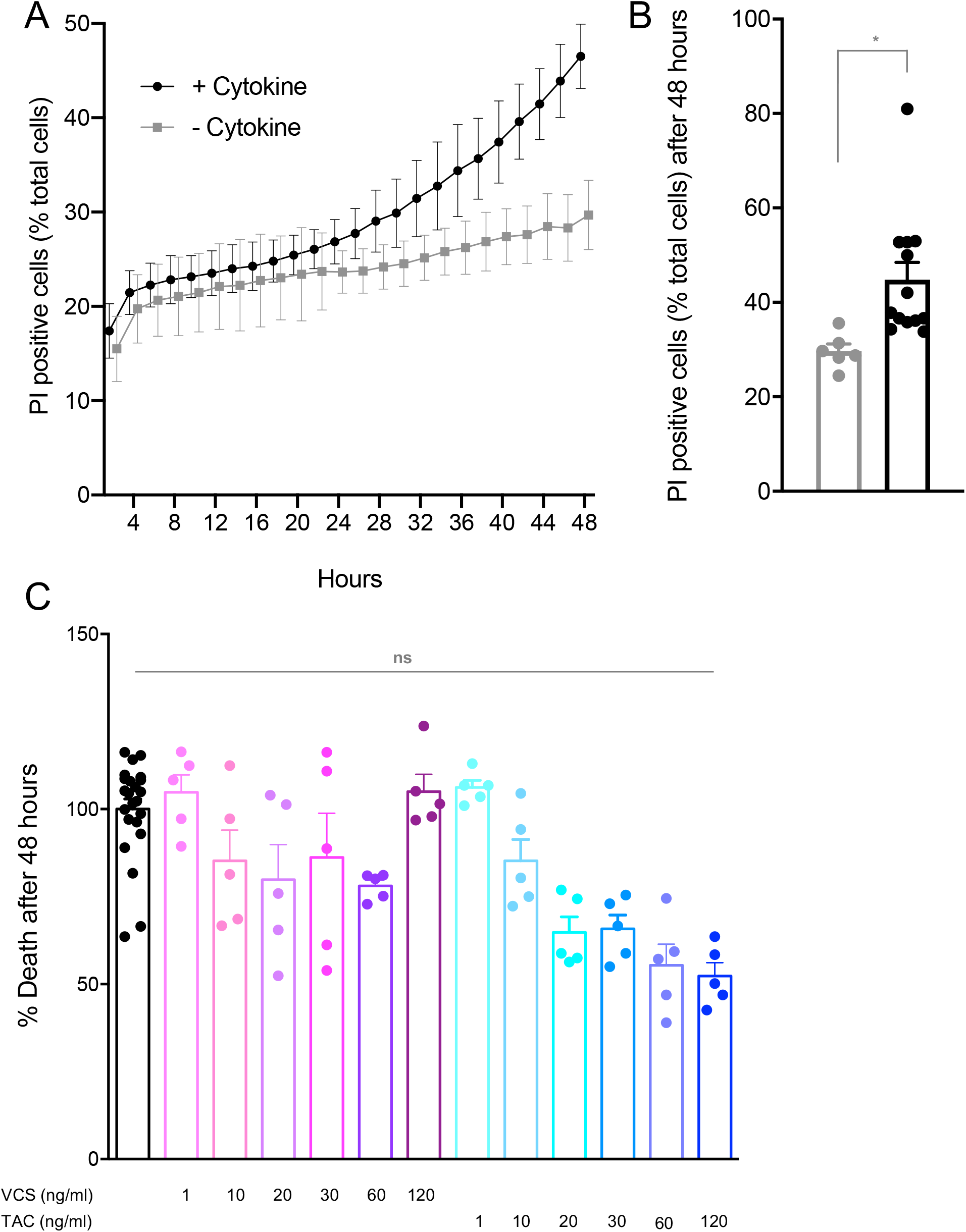
Islet cell survival from dispersed human islets treated with VCS and TAC. **(A,B)** Cell death was assessed every 30 minutes over 48 hours using Propidium Iodide (PI) and Hoechst staining, with a significant increase seen in the presence of a β-cell toxic cytokine cocktail. **(C)** Shown is the maximum cell death normalized measurement (see methods) for each condition TAC and VCS dose, normalized to the DMSO control cultures. Results shown are quantified from 5 cultures per drug condition, over two separate runs.

### Effects of TAC or VCS on human islet gene expression

In order to provide clues as to the underlying molecular mechanisms by which TAC, but not VCS, significantly inhibits insulin secretion we performed RNAseq and network analysis on human islets from 7 donors (Fig 1B). Despite significant donor to donor variation in gene expression, we were able to identify multiple significantly differentially expressed genes in human islets treated with these immunosuppressant drugs, compared to DMSO (Fig. 6A). Specifically, TAC significantly decreased the expression of synaptotagmin 16 (*SYT16*), TBC1 domain family member 30 (*TBC1D30*), phosphoenolpyruvate carboxykinase 1 (*PCK1*), SPARC related modular calcium binding 1 (*SMOC1*), synaptotagmin 5 (*SYT5*), muscular LMNA interacting protein (*MLIP*), coiled-coil domain containing 184 (*CCDC184*), ankyrin repeat and SOCS box containing 4 (*ASB4*), pyruvate dehydrogenase kinase 4 (*PDK4*), long intergenic non-protein coding RNA 473 (*LINC00473*), long intergenic non-protein coding RNA 602 (*LINC00602*), SLC8A1 antisense RNA 1 (*SLC8A1-AS1*), leucine rich repeat LGI family member 2 (*LGI2*), islet amyloid polypeptide (*IAPP*), cAMP responsive element modulator (*CREM*), B9 domain containing 1 (*B9D1*), and scavenger receptor class B member 2 (*SCARB2*). VCS only significantly decreased the expression of *SYT16*, *TBC1D30*, and *PCK1*, and to a qualitatively lesser degree.

**Figure 6.**
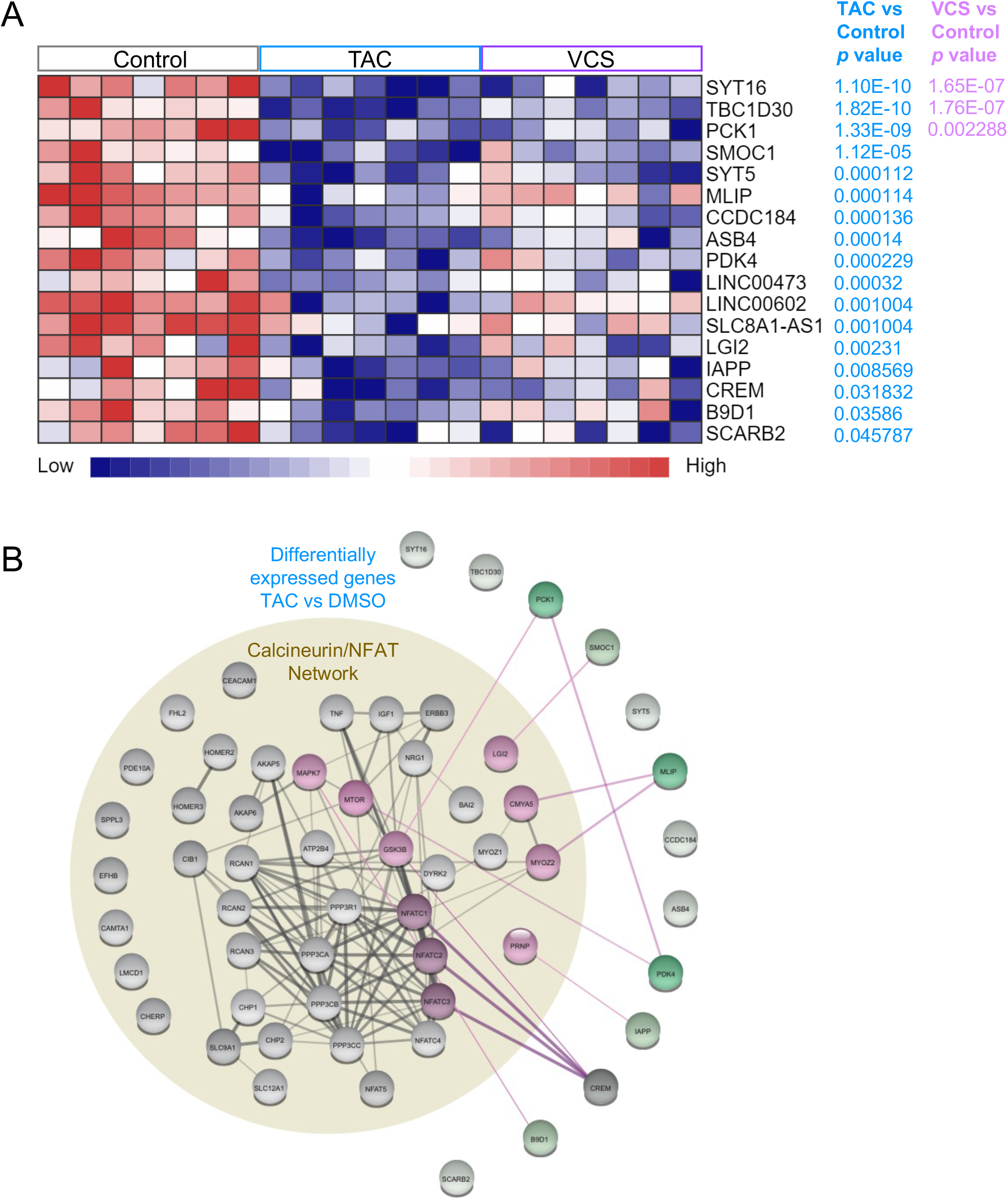
RNA sequencing analysis and protein-protein interaction networks of human islets treated with TAC and VCS at peak doses. **(A)** mRNAs that were nominally differentially expressed (adjusted *p* < 0.05) are shown **(B)** Protein-protein interaction networks highlight connections between calcineurin/NFAT pathway and deferentially expression genes in **(A)** VCS or **(B)** TAC treatment. Up or down regulated genes are shown in red or green, respectively.

Next, we examined the relationship between the calcineurin-NFAT gene network, as defined by its Gene Ontology term, and the TAC-modulated genes in human islets using single String protein-protein interaction network modelling (Fig. 6B). These analyses showed that CREM was the TAC-target mostly highly connected to calcineurin-NFAT network, with highest confidence-level interactions with NFATC1, NFATC2, and NFATC3, and a medium confidence link to GSK3B. MILP showed high confidence interactions with a module including Cardiomyopathy Associated 5 (CMYA5) and Myozenin 2 (MYOZ2). PCK1 showed a medium confidence interaction with GSK3B. PDK4 showed a high confidence interaction with MTOR, and also with PCK1.

We found medium confidence interactions between SMOC1 and Leucine Rich Repeat LGI Family Member 2 (LGI2), between B9D1 and Mitogen-Activated Protein Kinase 7 (MAPK7), and between IAPP and Prion Protein (PRNP). Together, these analyses show how VCS and TAC treatment affect the expression of specific genes via calcineurin signalling and lead us to our current working model (Fig. 7C).

**Figure 7.**
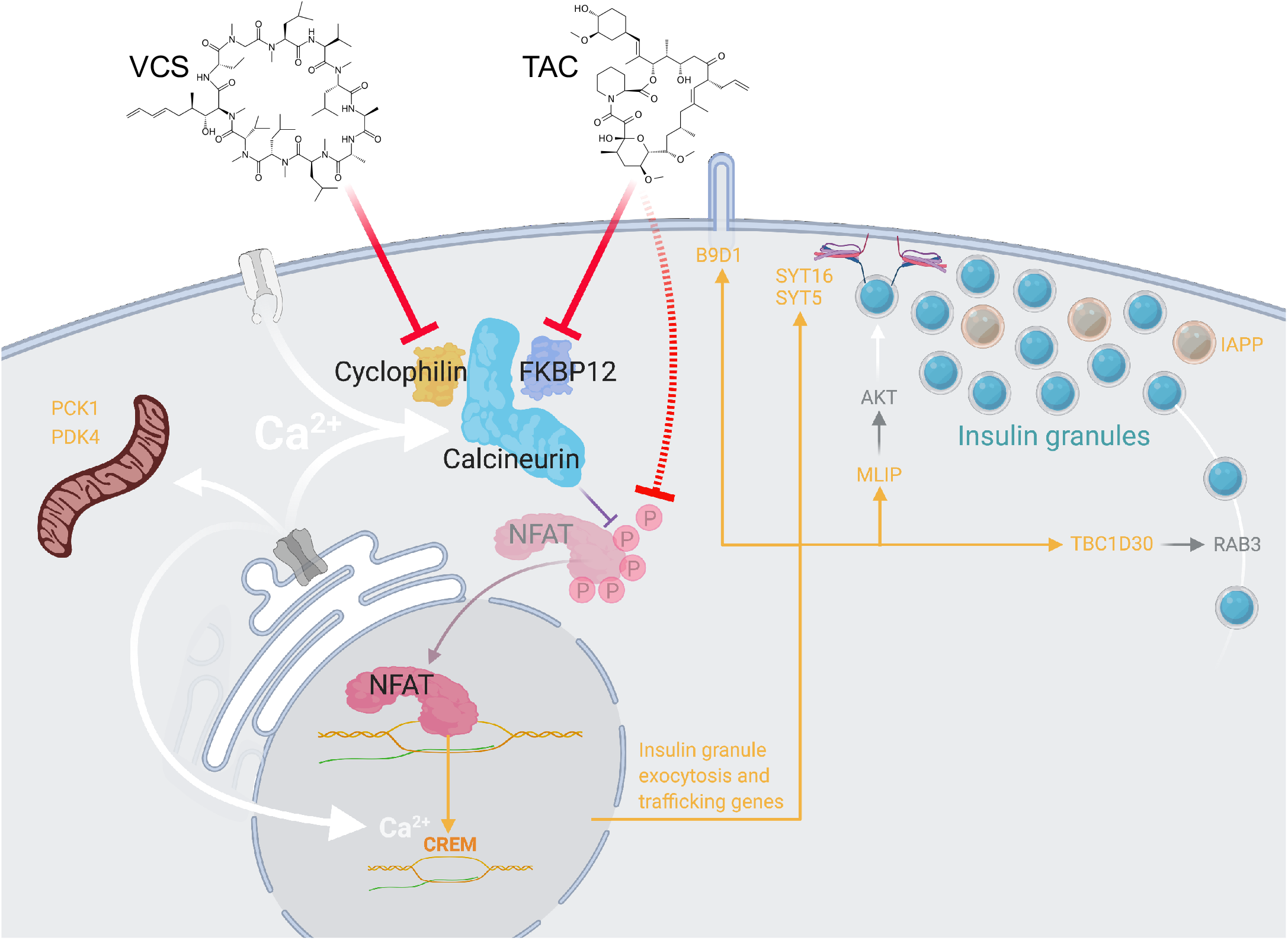
Working model of differential regulation of insulin exocytosis by VCS and TAC. In our model, TAC has more potent inhibitory effects on NFAT, and therefore results in the stronger suppression of key genes. TAC-sensitive gene products with known functions related to insulin secretion are shown in orange.

## Discussion

The effects of calcineurin inhibitors on human β-cells remain understudied. A deeper understanding of how immunosuppressive agents affect human islets is necessary to elucidate the pathogenesis of NODAT and to identify immunosuppressants with improved safety profiles. Thus, the objective of the present study was to determine why the next-generation calcineurin inhibitor, VCS, has lower human β-cell toxicity compared with TAC. To examine this, we compared the effects of TAC and VCS on the dynamics of insulin secretory function, the programmed cell death, and the transcriptomic profile of isolated human islets. We found that TAC, but not VCS, caused a significant impairment of both 15 mM glucose and 30 mM KCl-stimulated insulin secretion at clinically relevant doses. We saw no effect on total insulin content or human islet cell survival with either drug. Gene pathway enrichment analysis of RNAseq data showed that TAC decreased the expression of genes that regulate exocytosis and genes that regulate transport from the ER to the Golgi apparatus. VCS treatment also induced these changes; however, it did so to a lesser extent. Thus, this greater inhibition of insulin secretion could provide an explanation of why the risk of NODAT is higher in patients taking TAC than the next generation calcineurin inhibitor VCS. The equivalently immunosuppressive dose ranges of VCS and TAC we used have dramatically different potencies on NFAT, suggesting that NFAT-mediated gene expression mediates the observed effects on exocytosis regulating gene expression.

Previous studies have found that calcineurin inhibitors impair glucose-stimulated insulin secretion in rodent islets and rodent cell lines (8,28–39), although through mechanisms that we show here are not validated in human islets. For example, we do not see evidence for a robust effect of TAC or VCS on insulin production or gene expression. Moreover, the majority of rodent studies (8,28–39) and previous studies using human islets (7,10,40–47) used concentrations of calcineurin inhibitors that are higher than seen clinically. Our study extends these previous findings by using clinically relevant doses of TAC and VCS, rigorously examining the kinetics of insulin release in human islets and by implicating a direct effect on insulin granule trafficking. TAC at 10 ng/ml (clinical trough) and 20 ng/ml (clinical peak) significantly inhibited total insulin secretion in response to 15 mM glucose and this inhibition was directly due to a specific effect on the 2^nd^ phase of insulin release. Furthermore, KCl-stimulated insulin secretion was decreased to a similar degree, implicating secretory pathways distal to Ca^2+^ entry via voltage-gated Ca^2+^ channels. Together, these data suggest that the readily releasable pool of insulin granules is not robustly affected by TAC-mediated calcineurin inhibition; instead, TAC most likely impairs pathways associated with insulin granule trafficking to the plasma membrane. While we did not directly look at exocytotic events in this study, the results of our RNAseq analysis are consistent with a TAC-mediated effect on insulin granule traffic and release. TAC has previously been suggested to impair glucose-stimulated insulin secretion in rat islets via a mechanism involving insulin granule exocytosis (39). Future studies using electrophysiology and imaging to examine insulin granule docking, priming and release will provide a more detailed biophysical explanation of TAC effects in human β-cells.

Our RNAseq data provided novel insights into the molecular mechanisms controlling insulin secretion from human β-cells, the effects of these drugs, and impetus for follow-up studies on several gene products. A prime candidate is SYT16, which has not previously been directly implicated in glucose-stimulated insulin secretion but that is known to be glucose responsive in human islets (48) and under circadian control in mouse islets (49). The causal role of SYT5 in insulin secretion remains unclear in primary human β-cells (50), but should also be investigated further. TAC likely also has direct effects on insulin granule trafficking given that TBC1D30 is an activating GTPase of RAB3 (51). MLIP activates AKT, which can promote insulin secretion (52), and is regulated by CREB/CREM transcription factors. Indeed, TAC significantly inhibited CREM, a member of the CREB family of transcription factors known as master regulators of β-cell function (53). Previous studies in a β-cell line have shown that TAC inhibits CREB transcriptional activity (54). *B9D1* contributes to the biogenesis of the primary cilium, which is required for normal insulin secretion (55). *PCK1* and *PDK4* regulate cellular metabolism, but require additional studies in human islets. TAC reduced *IAPP* expression but the functional consequence of this is unclear. Thus, the majority of significantly downregulated genes point to the distal stages of insulin trafficking and exocytosis as the primary sites of TAC-induced β-cell dysfunction. We used protein-protein interaction mapping to predict the most likely link(s) between calcineurin signaling and the TAC-downregulated genes. Indeed, this network analysis suggested direct and robust links to CREM in TAC-treated human islets. Collectively, these data further solidify the role of calcineurin signaling and NFAT-mediated gene transcription in human β-cell function and provide insight into the molecular mechanisms controlling 2^nd^ phase insulin exocytosis.

Calcineurin activity has been ascribed a number of functions in pancreatic β-cells (8–10,41,56–59). For example, cultured insulinoma cells treated with the calcineurin inhibitor cypermethrin exhibited decreased insulin exocytosis in response to both glucose and KCl stimulation (60). In contrast to this, some studies suggest that calcineurin has a negative role in insulin exocytosis. However, these were indirect findings obtained by examining the inhibition of β-cell exocytosis via neurotransmitter-mediated activation of calcineurin (61,62), as opposed to direct effects observed following inhibition of calcineurin. Nevertheless, our study now shows that VCS is gentler on human islets because it does not cause the same level of inhibition on insulin exocytosis compared to TAC.

In our previous work, we examined the different effects of immunosuppressive drugs (TAC, cyclosporine, and rapamycin) on induced endoplasmic reticulum (ER) stress and caspase-3-dependent apoptosis in human islets (7). In that study, we found that TAC impaired glucose-stimulated insulin release from human islets, but did not significantly elevate the protein levels of CHOP or cleaved caspase-3, indicating that it’s effects were mostly on β-cell function rather than fate (7). In our current study we measured human islet cell survival over a longer period by using our high-content imaging protocol. We found that neither TAC nor VCS had significant deleterious effect on islet cell survival or insulin content over 48 hours, under the current testing conditions. Similarly, our post RNA-seq pathway analysis did not identify significant changes in genes involved in β-cell apoptosis and/or survival. While an increase in β-cell ER-stress and apoptosis is a plausible pathophysiological mechanism for the diabetic effects of immunosuppressants in the post-transplant clinical setting, it is unlikely that this is the mechanism explaining why the risk of NODAT is higher in patients taking TAC when compared with VCS.

Limitations of our study include the following. Although we had the benefit of directly examining human islets, our *in vitro* experiments may be impacted by islet isolation and culture. Nevertheless, the fundamental *in vivo* mechanisms appear to be well conserved in high quality isolated islets such as those used in our study (11). Mechanistically, our analysis of insulin secretion in response to high glucose and high KCl, together with the transcriptomic profiling, points to direct effects on the distal processes of insulin exocytosis. Future studies could directly measure exocytosis with electrophysiological capacitance measurements. Our data point to differences in NFAT activation between VCS and TAC than can explain the distinct impacts on insulin secretion, but our work employed non-β-cells for the dose-finding experiment and ideally future studies will be repeated in human β-cells or human β-cell lines. We chose the concentrations for our *in vitro* studies based on clinical efficacy. The observation that lower concentrations of TAC were more toxic than VCS favours this next generation immunosuppressant. Given the clinical importance of calcineurin inhibitor mediated β-cell dysfunction, we hope that our work stimulates more experimentation in human islets and other key tissue governing glucose homeostasis.

In summary, we have demonstrated that TAC, but not VCS, significantly inhibits the total response of glucose-stimulated insulin secretion from human islets. This decrease in secretion is not mediated by effects on islet cell health or insulin content and is most likely a direct consequence of decreased insulin granule exocytosis. VCS, a next generation calcineurin inhibitor, does not cause the same degree of inhibition of insulin secretion. Thus, this is a plausible physiological mechanism explaining the lower incidence of NODAT in patients taking VCS.

## Acknowledgments

Human islets for research were provided by the Alberta Diabetes Institute IsletCore at the University of Alberta in Edmonton (http://bcell.org/human-islets) with the assistance of the Human Organ Procurement and Exchange (HOPE) program, Trillium Gift of Life Network (TGLN) and other Canadian organ procurement organizations. Islet isolation was approved by the Human Research Ethics Board at the University of Alberta (Pro00013094). All donors’ families gave informed consent for the use of pancreatic tissue in research and we are grateful for their gift. We thank Xioake Betty Hu for technical assistance.

## Author Contributions

JK designed, conducted and analyzed experiments, and co-wrote the manuscript.

LB conducted and analyzed experiments and edited the manuscript.

PO designed, conducted and analyzed experiments, and edited the manuscript.

HHC designed and analyzed experiments and edited the manuscript.

EP designed, conducted and analyzed experiments, and edited the manuscript.

DRU designed, conducted and analyzed experiments, and edited the manuscript.

JLC designed experiments and edited the manuscript.

RBH designed experiments and edited the manuscript.

JDJ designed and analyzed experiments and co-wrote the manuscript.

## Data Availability

All data generated or analyzed during this study are included in this published article or in the data repositories listed in References.

